# Leveraging uncertainty quantification to optimise CRISPR guide RNA selection

**DOI:** 10.1101/2024.02.01.578527

**Authors:** Carl Schmitz, Jacob Bradford, Robert Salomone, Dimitri Perrin

## Abstract

CRISPR-based genome editing relies on guide RNA sequences to target specific regions of interest. A large number of methods have been developed to predict how efficient different guides are at inducing indels. As more experimental data becomes available, methods based on machine learning have become more prominent. Here, we explore whether quantifying the uncertainty around these predictions can be used to design better guide selection strategies. We demonstrate how using a deep ensemble approach achieves better performance than utilising a single model. This approach can also provide uncertainty quantification. This allows to design, for the first time, strategies that consider uncertainty in guide RNA selection. These strategies achieve precision over 91% and can identify suitable guides for more than 93% of genes in the mouse genome. Our deep ensemble model is available at https://github.com/bmdslab/CRISPR_DeepEnsemble.

## Introduction

CRISPR-based methodologies have established themselves as a very important instrument for genomic manipulation (1). Fundamentally, these technologies employ a CRISPR-associated (Cas) endonuclease alongside a sequence-specific RNA component that directs the nuclease towards a predetermined genomic locus. This guide RNA (gRNA) is engineered to target specific genomic sequences for editing. Specifically, in the context of Cas9, a potential CRISPR target site requires the presence of a protospacer adjacent motif (PAM) characterized by an NGG sequence, with the adjacent 20 nucleotide sequence upstream serving as a template for constructing the gRNA.

Over the preceding decade, the scientific community has leveraged CRISPR technology for a wide array of purposes, ranging from foundational research to practical applications. These include the development of animal models for disease research (2), the genetic study of endangered species (3), enhancement of agricultural crop resilience (4), and the pioneering of novel therapeutic approaches (5).

However, despite the broad spectrum of applications underscoring the versatility of CRISPR-based genome editing, the process of gRNA design remains a non-trivial and intricate task, demanding careful consideration and expertise.

One of the objectives when designing gRNAs is to maximise the on-target efficiency, which can be understood as the rate at which the desired edit is obtained. Liu et al. (6) highlights how a wide range of factors have been investigated to determine the effects on gRNA efficiency. To assist in the process, many tools have been developed (7, 8).

While most of the tools can identify some efficient guides, the overlap between them is often limited (7). While this behaviour can be exploited to develop consensus approaches that outperform individual tools (9, 10), there exists considerable room for further improvements.

As more experimental data have become available, improvements have increasingly been sought through machine learning approaches, with a particular focus on deep learning (11– 14).

One limitation of complex machine learning methods is the risk of treating them as black boxes and putting too much trust in their output. This is particularly true for deep learning, where explainability is a challenge.

In this paper, we explore the notion of uncertainty. Can we deploy simple and scalable strategies to estimate the uncertainty in the predicted efficiency? If this is achievable, can we develop new guide RNA design strategies that incorporate that uncertainty, and will that improve the quality of the guides being selected?

## Materials and Methods

### A. CRISPRon

Xiang et al. (14) experimentally generated indel frequencies of 10,592 Cas9 gRNAs with minimal overlap with existing datasets that also report indel frequencies. Using this data, they developed a deep learning model called CRISPRon. The initial CRISPRon model was trained using one-hot encodings of 30 bp sequences (4 + spacer 20 + PAM 3 + 3) and other features such as melting point, and RNA-DNA binding energy. The initial results showed a strong correlation with the existing dataset from (12). This resulted in the merging of the two datasets to create a new dataset of 23,902 guides. The merged dataset was used to train the final version of CRISPRon. CRISPRon was reported to have a better correlation on both an internal independent test set and an external test set, outperforming popular models such as as Azimuth (11), DeepSpCas9 (12) and DeepHF (13).

The CRISPRon model is essentially a convolutional neural network taking the 30bp sequence as input, modified to take additional features as input to its later layers, which resemble those of a regular feedforward neural network. See Xiang et al. (14, Figure 2.a) for a diagrammatic representation of the precise neural network architecture.

**Fig. 1.**
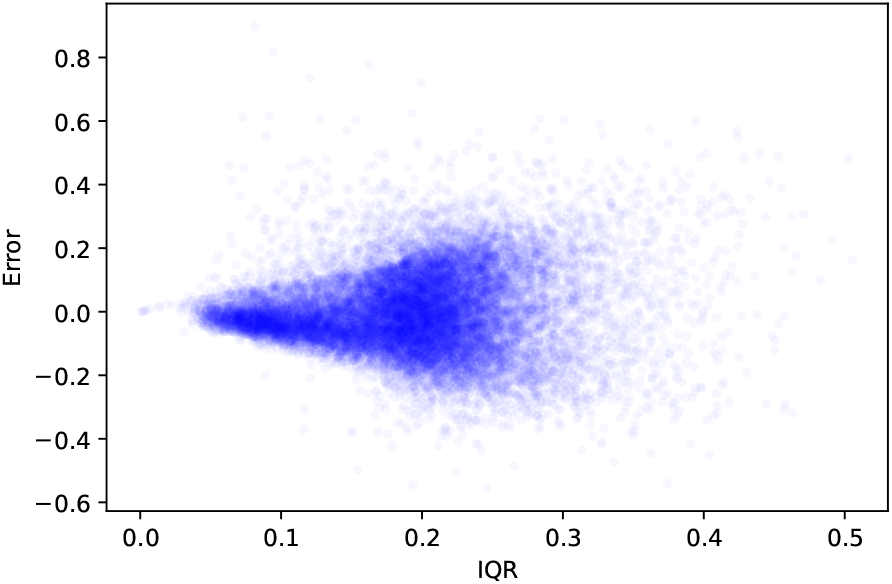
Absolute prediction error as function of uncertainty (measured using IQR)

**Fig. 2.**
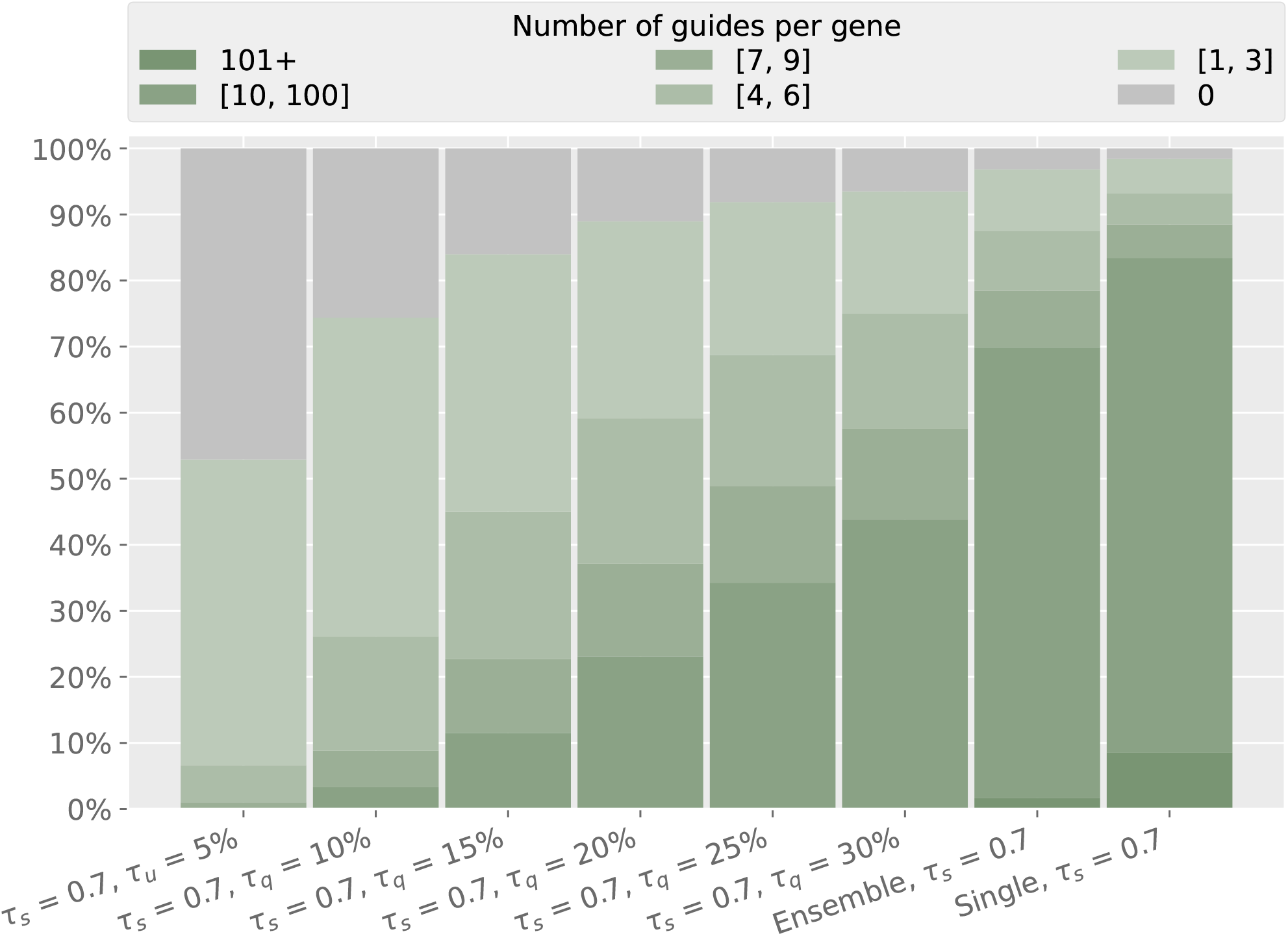
Accepted guides per gene (binned by guide counts)

Given the strong performance of the model, we implemented it in PyTorch (15) as a base model which we modified to allow for uncertainty quantification.

### B. Uncertainty quantification and deep ensemble approach

We modify the CrisprON model in two ways, to capture both the aleotoric (data-based) and epistemic (modelbased) uncertainties in the predictions.

For the former, rather than outputting a single prediction, the inherent variability in the data is accounted for by modelling the response variable as coming from a Beta distribution (which takes values between zero and one, the range of possible efficiency values), whose parameters are given by the output of the neural network. That is, the neural network outputs a *distribution* as a prediction implicitly via giving the two parameters of a Beta distribution (which will be different for each input). The model is trained via maximum-likelihood, that is, choosing the parameters that maximize the average likelihood over the observations for the output Beta distributions. A prediction can be obtained by outputting the expected value of the respective Beta distribution. The uncertainties for a single model can be obtained by simulating many times from the corresponding Beta distribution.

Whilst the above approach is capable of modelling the uncertainty in the response variable for a given input, it does not capture our inherent uncertainty in the model itself that should be used to make such predictions.

To overcome the latter issue, we use a simple *deep ensemble* approach. The essence of the approach is simple — one trains not one model but instead some large number of models, each with a different initialisation for the training procedure. This results in a collection of different models, due to the existence of many local minima in the training objective function (different initialisation will result in the training ending up in different minima).

Despite the apparent simplicity, such an approach is effective as deep ensembling can be viewed as a crude approximation to sampling from the posterior distribution of a Bayesian model (16), which is a standard approach for accounting for model-based uncertainty (but is typically computationally intractable in deep learning settings). An additional advantage of deep ensembles is that they tend to produce better results in terms of performance (generalization to unseen data) (17).

As mentioned, our initial modelling uses a probabilistic (Beta distribution based) extension of (14). However, instead of using a single model, we fit an ensemble of 25 models. Each ensemble member was trained for 50 epochs. The input for our deep-learning model uses the same 30 bp sequence (4 + spacer + PAM + 3) and uses the sequence melting point as a feature.

The ensemble approach is configured as an unweighted model average, where all of the predictions for the ensemble members are averaged to give a final prediction.

Uncertainty bands for prescribed quantiles can be produced by simulating many times from each individual ensemble members Beta distribution, aggregating the resulting samples, and computing the empirical quantiles of this simulated response.

### C. Data

We combined the CRISPRon (14) datasets with other datasets containing experimental results on indel frequency (12, 13, 18). All datasets were filtered to remove duplicated entries and NaN results.

The datasets (13, 14) were expanded to 30 bp sequences. The expansion was achieved by aligning the provided sequences to the reference genome using Bowtie2 (19), and extracting the extended sequences with SAMtools (20) to extract the extended sequences. Any sequence that had multiple perfect alignments from Bowtie2 was excluded.

After the above preprocessing, 13,359 guides remained from (12), 49,523 from (13), 8,372 from (18) and 9,886 from (14), yielding 80,408 guides in total. The processed datasets were then merged into a final dataset, and guides that were present in multiple datasets were merged and averaged.

The 30 bp sequences were individually one-hot encoded and their respective melting points were calculated. Finally, our processed dataset was divided into training and testing portions, of sizes 60,408 and 20,000 guides, respectively.

### D. Metrics and thresholds

Three methods are used to evaluate the performance of approaches involving our ensembled model and its associated uncertainty ranges.

The first method compares the *predicted* score from the model with the observed indel frequency, i.e., the *actual* score. This considers the correlation between the scores, as in (14), and the absolute prediction error.

To explore guide design strategies, it is convenient to transform the scores into binary classes. Is this guide efficient or not? Is it accepted by the model or not?

The second level of evaluation is then to consider the level of performance for this binary decision problem. For any gene, there are often dozens of guides that can be used, so it is more crucial that the selected guides are efficient, rather than to select *all* the efficient guides. As a result, precision is the key metric of interest. For completeness, recall is also reported.

The third level of evaluation tests this assumption that a high recall is not essential. All potential CRISPR sites in the mouse genome are extracted and evaluated. Then, we count the number of selected guides for each gene.

The selection of a threshold to turn the observed indel frequencies into binary classes is somewhat arbitrary. Based on the distribution of the training and on considerations on practical constraints of genome editing experiments, we chose 0.7 as the default. To explore the impact of that choice, we also repeated all tests for 0.6 and 0.8.

To make the binary decision of selecting a guide or not, we can consider the score alone, or the score in conjunction with some notion of uncertainty (difference between lower and upper bound in the predictions, or interquartile range). In what follows, we note *τ*_*s*_ the threshold on the score, *τ*_*u*_ the thresh-old on the difference between the bounds, and *τ*_*q*_ the thresh-old on the interquartile range. For *τ*_*s*_, the threshold is defined directly on the predicted score (i.e. single score if using a single model, or average score if using an ensemble). For *τ*_*u*_ and *τ*_*q*_, the threshold is defined based on the range of values observed with the training data. For instance, *τ*_*q*_ = 20% means we pick *τ*_*q*_ so that 20% of the guides used for training had an interquartile range in predicted scores below *τ*_*q*_ (which corresponds to a threshold of 0.13). To select guides, we only choose guides that have a predicted score above *τ*_*s*_ *and* an uncertainty threshold (if used) below *τ*_*u*_ (or *τ*_*q*_).

These three levels of evaluation are used to assess our ensemble approach. To understand the contribution of the ensembling and uncertainty quantification, we also use a single prediction from the first model of the ensemble. Note that this single model should not be seen as an exact replication of CRISPRon, due to the use of the Beta distribution as described above.

## Results

### E. Scoring performance

After training the ensemble, the performance was assessed on the testing set of 20,000 guides. The Spearman correlation and Pearson correlation were 0.839 and 0.838, respectively. To understand the performance of the ensemble, the first member was selected and tested in isolation, resulting in a Spearman correlation and Pearson correlation of 0.706. The ensemble model provides superior performance.

Next, we looked at the absolute error in the predicted score. For the ensemble model, we observe a mean absolute error of 0.0947 (standard deviation 0.0813). In contrast, the single model had mean of 0.1395 and standard deviation of 0.1268. Again, there is a clear advantage to the deep ensemble.

Figure 1 shows the error as a function of uncertainty (measured using IQR). The lower the uncertainty, the lower the prediction error. This further highlights the benefits of the ensemble approach. It also provides a motivation for using that uncertainty in the guide selection: if the uncertainty is low, the predicted score is more likely to be accurate, and filtering for highly-scored guides should return efficient ones.

### F. Guide selection performance

Next, we explored different guide selection strategies to understand how the ensembled model and the uncertainty quantification could be exploited to select more efficient guides. We varied the threshold *τ*_*s*_ on the predicted score, and the thresholds *τ*_*u*_ and *τ*_*q*_ on the uncertainty metrics.

Table 1 shows some of the top-performing threshold combinations, sorted by precision. It is not a surprise to see a majority of configurations using *τ*_*s*_ = 0.7, given how we binarized the actual scores into efficient/inefficient classes: in the default configuration, we used a threshold of 0.7 on the actual score. We explore this in Section H.

**Table 1.**
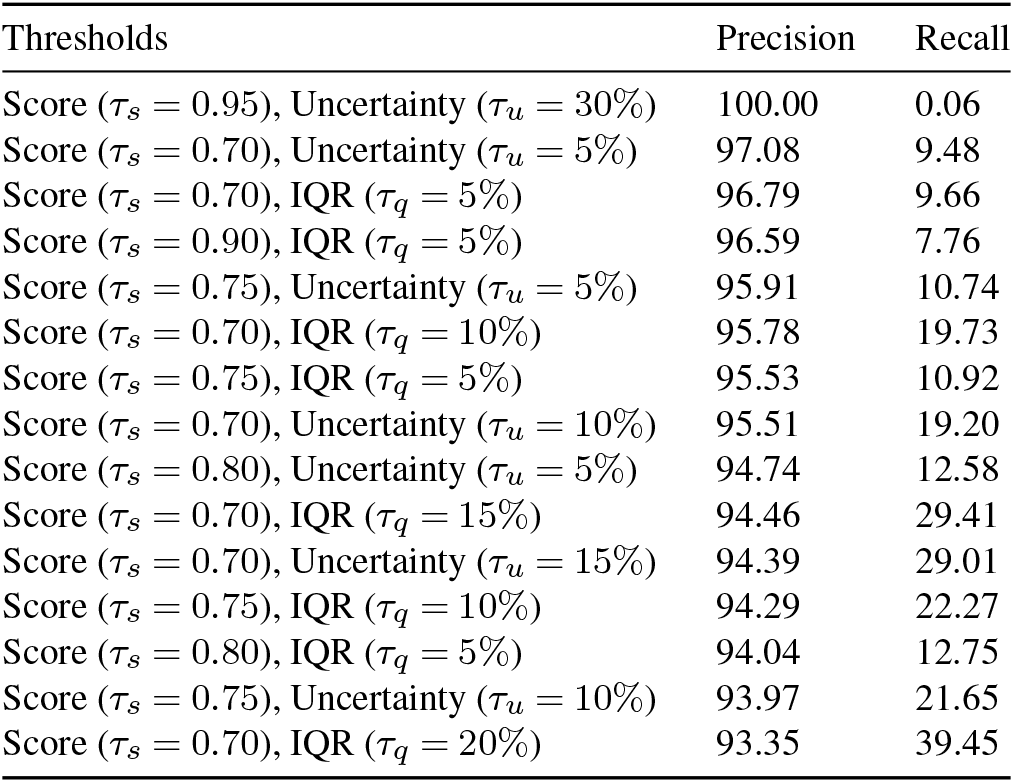
Some of the best configurations based on the 20K holdout testing set. Results are ordered by precision and duplicates are removed.

| Thresholds | Precision | Recall |
| --- | --- | --- |
| Score ($\tau_s = 0.95$), Uncertainty ($\tau_u = 30\%$) | 100.00 | 0.06 |
| Score ($\tau_s = 0.70$), Uncertainty ($\tau_u = 5\%$) | 97.08 | 9.48 |
| Score ($\tau_s = 0.70$), IQR ($\tau_q = 5\%$) | 96.79 | 9.66 |
| Score ($\tau_s = 0.90$), IQR ($\tau_q = 5\%$) | 96.59 | 7.76 |
| Score ($\tau_s = 0.75$), Uncertainty ($\tau_u = 5\%$) | 95.91 | 10.74 |
| Score ($\tau_s = 0.70$), IQR ($\tau_q = 10\%$) | 95.78 | 19.73 |
| Score ($\tau_s = 0.75$), IQR ($\tau_q = 5\%$) | 95.53 | 10.92 |
| Score ($\tau_s = 0.70$), Uncertainty ($\tau_u = 10\%$) | 95.51 | 19.20 |
| Score ($\tau_s = 0.80$), Uncertainty ($\tau_u = 5\%$) | 94.74 | 12.58 |
| Score ($\tau_s = 0.70$), IQR ($\tau_q = 15\%$) | 94.46 | 29.41 |
| Score ($\tau_s = 0.70$), Uncertainty ($\tau_u = 15\%$) | 94.39 | 29.01 |
| Score ($\tau_s = 0.75$), IQR ($\tau_q = 10\%$) | 94.29 | 22.27 |
| Score ($\tau_s = 0.80$), IQR ($\tau_q = 5\%$) | 94.04 | 12.75 |
| Score ($\tau_s = 0.75$), Uncertainty ($\tau_u = 10\%$) | 93.97 | 21.65 |
| Score ($\tau_s = 0.70$), IQR ($\tau_q = 20\%$) | 93.35 | 39.45 |

What is more interesting is that we obtain very high precision, with all these configurations scoring above 93%. The guides being selected have a very high chance of being efficient in practice.

It is also interesting to note that the difference between using a threshold on the whole uncertainty range (*τ*_*u*_) or on the interquartile range (*τ*_*q*_) does not make a big difference. The threshold value is more important than its scope.

As expected, the recall values are lower. This is a direct consequence of prioritising precision. The assumption is that these recall values (especially those above 15%) are high enough to select enough guides for most genes. We explore this in Section I.

When the uncertainty threshold is too low, *τ*_*s*_ becomes redundant. We do not see predictions where the uncertainty is very low and the predicted score is low, so a very tight constraint on the uncertainty means that only high-score predictions are selected. Similarly, if the score threshold is extremely high, the uncertainty is always low (because the prediction is an average and is bounded by 1, so it can only be very high if all individual scores are high) and the uncertainty threshold becomes redundant. This leads to identical sets of results, so Table 1 does not show duplicates. These extreme configurations, such as *τ*_*s*_ = 0.7, *τ*_*q*_ = 5% or *τ*_*s*_ = 0.95, *τ*_*q*_ = 30%, are not considered practical.

In Table 2, we fix the score threshold at *τ*_*s*_ = 0.7, and explore the impact of the deep ensemble and of uncertainty quantification. Just using the ensemble offers a 9 percentage points increase in precision (75.17 to 84.27%). Taking the uncertainty into account add another 6 to 13 percentage points in precision. Of course, the cost is a lower recall, but some configurations, such as *τ*_*s*_ = 0.7, *τ*_*q*_ = 30% still provide a recall above 55%.

**Table 2.** Impact of the uncertainty threshold. All configurations use a score threshold of 0.7. Results are ordered by precision.

| Thresholds | Precision | Recall |
| --- | --- | --- |
| Score ($\tau_s = 0.70$), Uncertainty ($\tau_u = 5\%$) | 97.08 | 9.48 |
| Score ($\tau_s = 0.70$), IQR ($\tau_q = 5\%$) | 96.79 | 9.66 |
| Score ($\tau_s = 0.70$), IQR ($\tau_q = 10\%$) | 95.78 | 19.73 |
| Score ($\tau_s = 0.70$), Uncertainty ($\tau_u = 10\%$) | 95.51 | 19.20 |
| Score ($\tau_s = 0.70$), IQR ($\tau_q = 15\%$) | 94.46 | 29.41 |
| Score ($\tau_s = 0.70$), Uncertainty ($\tau_u = 15\%$) | 94.39 | 29.01 |
| Score ($\tau_s = 0.70$), IQR ($\tau_q = 20\%$) | 93.35 | 39.45 |
| Score ($\tau_s = 0.70$), Uncertainty ($\tau_u = 20\%$) | 92.82 | 38.13 |
| Score ($\tau_s = 0.70$), IQR ($\tau_q = 25\%$) | 92.48 | 48.34 |
| Score ($\tau_s = 0.70$), Uncertainty ($\tau_u = 25\%$) | 91.68 | 45.70 |
| Score ($\tau_s = 0.70$), IQR ($\tau_q = 30\%$) | 91.11 | 55.96 |
| Score ($\tau_s = 0.70$), Uncertainty ($\tau_u = 30\%$) | 90.74 | 51.97 |
| Ensemble, Score ($\tau_s = 0.70$) | 84.27 | 80.33 |
| Single Model, Score ($\tau_s = 0.70$) | 75.17 | 76.97 |

### G. Previous datasets

In order to further our understanding of the model performance, we used the dataset generated by (21). We took all guides from this dataset and removed those that appeared in our training data. We ran different configurations of our model on the remaining guides. The results are shown in Table 3.

**Table 3.** Selected threshold configurations tested on the filtered Wang dataset, compared with Crackling.

| Thresholds | Precision | Recall |
| --- | --- | --- |
| Score ($\tau_s = 0.70$), Uncertainty ($\tau_u = 5\%$) | 100.00 | 0.98 |
| Score ($\tau_s = 0.70$), IQR ($\tau_q = 10\%$) | 100.00 | 2.66 |
| Score ($\tau_s = 0.70$), IQR ($\tau_q = 30\%$) | 95.92 | 13.17 |
| Score ($\tau_s = 0.70$), IQR ($\tau_q = 25\%$) | 94.81 | 10.22 |
| Score ($\tau_s = 0.70$), IQR ($\tau_q = 15\%$) | 94.44 | 4.76 |
| Score ($\tau_s = 0.70$), IQR ($\tau_q = 20\%$) | 94.44 | 7.14 |
| Crackling ($N = 3$) | 89.91 | 13.73 |
| Crackling ($N = 2$) | 83.82 | 44.26 |

It is important to note that for this dataset the efficient/inefficient classes are based on the *log*_2_ fold change, not indel frequency. Some indels may not lead to a change in expression, so it is a more difficult task.

For all configurations, we obtained a high precision, ranging from 94.44% to 100%. This is higher than the results obtained by Crackling, at the cost of a lower recall.

On the complete dataset, Crackling was reported to outperform all the tools it was tested against (10), so these results are encouraging.

### H. Impact of threshold choice

As discussed in Section D, the threshold used to transform the indel frequency into binary classes is somewhat arbitrary. To ensure that the results described in the previous Section are not an artefact of that choice, we repeated the same evaluation with a lower and a higher threshold. Overall, the results are consistent:

Precision is very high. Recall is lower, but for many threshold configurations it is sufficiently high.

Extreme threshold values for either score or uncertainty make the other threshold redundant and produce a very narrow set of guides (with high precision but a very low recall).

A threshold score close to the threshold to define efficiency generally produces good performance, but other configurations can also work.

Table 4 shows the impact of the efficiency threshold on a range of configurations. Our model reaches a high precision for all configurations if the efficiency boundary is at an indel frequency of 0.6 or 0.7. A boundary at 0.8 is more challenging, but several configurations still reach a precision above 91%. These results confirm that the model generalises well.

**Table 4.** Impact of the threshold used to define efficiency.

| Thresholds | Efficiency at 0.6 |  | Efficiency at 0.7 |  | Efficiency at 0.8 |  |
| --- | --- | --- | --- | --- | --- | --- |
|  | Precision | Recall | Precision | Recall | Precision | Recall |
| Score ($\tau_s = 0.60$), IQR ($\tau_q = 10\%$) | 97.34 | 16.27 | 95.78 | 19.73 | 91.90 | 25.73 |
| Score ($\tau_s = 0.60$), IQR ($\tau_q = 15\%$) | 96.40 | 24.35 | 94.46 | 29.41 | 88.84 | 37.60 |
| Score ($\tau_s = 0.60$), IQR ($\tau_q = 20\%$) | 96.00 | 32.90 | 93.35 | 39.45 | 86.02 | 49.40 |
| Score ($\tau_s = 0.60$), IQR ($\tau_q = 25\%$) | 95.54 | 40.55 | 92.40 | 48.35 | 83.21 | 59.17 |
| Score ($\tau_s = 0.60$), IQR ($\tau_q = 30\%$) | 94.97 | 47.61 | 90.81 | 56.13 | 79.68 | 66.94 |
| Score ($\tau_s = 0.70$), IQR ($\tau_q = 10\%$) | 97.34 | 16.27 | 95.78 | 19.73 | 91.90 | 25.73 |
| Score ($\tau_s = 0.70$), IQR ($\tau_q = 15\%$) | 96.40 | 24.35 | 94.46 | 29.41 | 88.84 | 37.60 |
| Score ($\tau_s = 0.70$), IQR ($\tau_q = 20\%$) | 96.00 | 32.90 | 93.35 | 39.45 | 86.02 | 49.40 |
| Score ($\tau_s = 0.70$), IQR ($\tau_q = 25\%$) | 95.58 | 40.52 | 92.48 | 48.34 | 83.30 | 59.17 |
| Score ($\tau_s = 0.70$), IQR ($\tau_q = 30\%$) | 95.11 | 47.38 | 91.11 | 55.96 | 80.09 | 66.86 |
| Score ($\tau_s = 0.80$), IQR ($\tau_q = 10\%$) | 97.34 | 16.27 | 95.78 | 19.73 | 91.90 | 25.73 |
| Score ($\tau_s = 0.80$), IQR ($\tau_q = 15\%$) | 96.40 | 24.33 | 94.45 | 29.39 | 88.84 | 37.57 |
| Score ($\tau_s = 0.80$), IQR ($\tau_q = 20\%$) | 96.05 | 32.47 | 93.56 | 39.00 | 86.43 | 48.97 |
| Score ($\tau_s = 0.80$), IQR ($\tau_q = 25\%$) | 95.80 | 38.23 | 93.15 | 45.83 | 85.00 | 56.84 |
| Score ($\tau_s = 0.80$), IQR ($\tau_q = 30\%$) | 95.58 | 41.47 | 92.76 | 49.63 | 84.30 | 61.30 |

### I. Whole-genome performance

Finally, we evaluated the performance of our model across the entire mouse genome. Here, we do not have a ground truth for all potential CRISPR sites. Instead, we use some of the best-performing methods from Section F, for which we know the precision, and assess whether their recall is sufficient to identify efficient guides across a large number of genes.

The results are shown in Figure 2. While some settings are too extreme (as previously discussed), the *τ*_*s*_ = 0.7, *τ*_*q*_ = 30% configuration performs very well. It can identify at least one guide for 93.50% of the genes, and our earlier evaluation had its precision at above 91%. For 80.99% of the genes, it can identify more than 3 guides. This enables multi-targeting, which has been shown to dramatically increase knockout efficiency (22).

Overall, this confirms that the deep ensemble approach can be used to identify guides for most genes that are close to being guaranteed to be efficient.

## Discussion

### J. Uncertainty can be quantified, and it can be used to improve guide selection

We proposed and tested a deep ensemble approach, and showed that it is able to outperform a single model using the same architecture.

Crucially, our deep ensemble also provides a method to quantify the uncertainty in the score prediction. We believe that it is the first method to do so in the context of CRISPR guide RNA design.

This uncertainty allows us to design novel guide selection strategies, which rely not only on the predicted score but also on how confident we can be in that score. We showed that these novel strategies achieve very high precision, and we confirm our hypothesis that while the recall is lowered, it remains high enough to identify guides for most genes.

The deep ensemble generalises well, including to datasets where guide efficiency is reported using different metrics.

### K. Future directions

This paper represents a first attempt at leveraging uncertainty quantification to design guide RNAs. While the results are very promising, there are a number of directions to explore.

Here, we used a Beta distribution to capture aleoteric uncertainty, but the approach can accommodate other distributions.

It would also be interesting to evaluate the impact of the size of the ensemble.

Methods based on the consensus between multiple approaches can produce good results (10). Combining this consensus philosophy with the ability to estimate uncertainty could lead to new solutions with improved recall.

As more experimental data continues to become available, the ensemble can be retrained and improved, or extended to other Cas proteins.

## Conclusions

In this paper, we investigated the use of deep ensembles to improve the prediction of the on-target efficiency of CRISPR guide RNAs. We showed that this approach can capture both the aleotoric and epistemic uncertainties in the predictions. We also showed that the ensemble provides a more accurate score and that, by combining it with the uncertainty estimates, we can design guide selection strategies with a very high precision. This comes with a lower recall, but we also showed that it remains high enough to identify suitable guide RNAs for most genes.

This represents a first attempt to leverage uncertainty quantification in CRISPR guide RNA design, and opens interesting directions for future research.

Our deep ensemble model is available at https://github.com/bmds-lab/CRISPR_DeepEnsemble.

## ACKNOWLEDGEMENTS

C.S. is supported by an Australian Government Research Training Program Scholarship. D.P. is supported by the Australian Research Council (ARC Discovery Project DP210103401).

